# Dual resistance to cold and ethanol in *Penicillium* species persistently contaminating donated cadavers

**DOI:** 10.1101/2025.11.10.687738

**Authors:** Tatsuya Yoshinouchi, Tomofumi Nakamura, Masahiro Manabe, Daisuke Mori, Jun-ichirou Yasunaga, Yasuhito Tanaka, Joe Iwanaga, Norio Kitagawa

**Author notes:** To whom correspondence should be addressed; Tomofumi Nakamura, M.D., Ph.D. (T.N.).

## Abstract

Modern cadaver preservation techniques, such as low-concentration formaldehyde methods, are increasingly adopted to mitigate health risks and improve tissue fidelity for medical education and surgical training. However, these environments potentially foster fungal contamination. This study aimed to identify contaminating species from cadaveric surfaces stored under specific preservation conditions and establish effective countermeasures.

Genetic identification revealed *Penicillium paneum* as the predominant species, while *Penicillium corylophilum* was also detected. These species exhibited distinctive traits, including vigorous growth at 4°C and marked ethanol tolerance, that explain their persistence in this preservation environment. Antifungal susceptibility testing demonstrated that these isolates were highly susceptible to commonly used disinfectants such as benzalkonium chloride, chlorhexidine gluconate, and phenol.

Our findings indicate that these *Penicillium species* are uniquely adapted to low-temperature and ethanol-rich preservation environments. The incorporation of the identified disinfectants into preservation solutions offers a practical and effective strategy for controlling persistent fungal contamination in cadaveric facilities.

## 1. Introduction

Fungal contamination of preserved cadavers poses a significant challenge in anatomical education facilities, potentially compromising both educational quality and occupational safety. While traditional high-concentration formalin methods effectively prevent microbial growth, health concerns have driven a transition toward safer preservation techniques with reduced formaldehyde exposure. However, these modified approaches may create environments conducive to fungal proliferation. Formaldehyde solution (formalin) was first documented for human cadaver embalming in 1899. Over a century later, the fundamental chemistry and techniques of formalin preservation remain largely unchanged (Musiał et al., 2016; Brenner et al., 2014).

The formalin method has been the predominant technique for preserving donated bodies, which are indispensable for medical education and research in Japan. This method, utilizing a fixative solution primarily composed of high-concentration formaldehyde, offers the significant advantage of long-term, stable preservation due to its potent bactericidal, fungicidal, and preservative properties. It has played a crucial role in effectively inhibiting tissue decomposition and reducing the risk of infection during anatomical dissection. However, the toxicity of formalin to human health has been a subject of significant concern (Chia et al.,1992). Exposure to formaldehyde is associated with a pungent odor, irritation of the eyes and respiratory tract, and dermatitis, while long-term exposure raises concerns regarding more severe health risks. Consequently, the need to protect the health of students and faculty has necessitated a shift toward safer preservation methods.

This has led to an accelerated transition at many institutions from high-concentration formalin-based fixation to alternative techniques, including the Thiel method which involves a reduced formalin concentration and other modified protocols (Eisma et al.,2013; Hayashi et al.,2014). The Thiel method, one of the prominent alternatives, employs a preservation solution composed primarily of substances such as ethanol and glycerin (Thiel et al., 1992). Cadavers preserved using the Thiel and similar modified methods exhibit remarkably life-like properties when compared to those fixed with conventional high-concentration formalin. Their tissues retain greater flexibility and joint movements are more natural, thereby providing a high-fidelity platform for practical medical training, including surgical simulations and endoscopic procedures (Hammer et al., 2022; Yiasemidou et al., 2017).

At Institute of Science Tokyo, we have adopted a modified preservation approach that combines 8% formalin perfusion with subsequent storage in 30% ethanol-soaked cloths at 4°C. This protocol aims to balance tissue preservation quality and reduced formaldehyde exposure. However, we recently observed fungal contamination on the cadaveric surfaces. This fungal overgrowth interfered with the gross anatomy laboratory, necessitating the removal of the fungi from the cadaveric surfaces prior to the sessions. The objective of this study was, therefore, to identify the fungal species capable of proliferating under these specific conditions and develop effective countermeasures to inhibit their growth on the surfaces of these donated bodies.

## 2. Material and Methods

### 2.1 Cadaver embalming and preservation protocol

Donated human cadavers were embalmed via femoral artery perfusion using 8% formalin solution without additional chemical agents. The preservation protocol consisted of two phases: Phase 1 (Pre-dissection): Cadavers were placed in Leichebags filled with 30% ethanol solution, sealed, and stored at 4°C in a cold storage room. Phase 2 (Post-dissection commencement): Cadavers were wrapped in towels soaked in 30% ethanol and stored at room temperature (approximately 20–25°C). Fungal contamination was first observed during Phase 1 storage on cutaneous surfaces of the chest, abdomen, hands, and legs.

### 2.2 Culture medium and fungal information

Sabouraud Dextrose Agar (SDA) medium was prepared with meat peptone (5 g/L), casein peptone (5 g/L), glucose (40 g/L), and agar (1.5% w/v). The medium was sterilized by autoclaving and subsequently supplemented with chloramphenicol (Cp) and kanamycin (K). Separately, RPMI 1640 medium (Nissui, Japan) was prepared without NaHCO_3_ and buffered to pH 6.8 with 0.165 M MOPS (3-morpholinopropane-1-sulfonic acid). This MOPS-buffered RPMI medium was then sterilized by filtration through a 0.25 µm filter and supplemented with Cp and K.

Representative isolates from this study, including two *Penicillium paneum isolates* (designated *P. paneum-*1 and *P. paneum*-2) and one *Penicillium corylophilum* isolate (*P. corylophilum*-1), were selected for further characterization. The following reference strains were used as controls: *Candida albicans* (NBRC-1594), *Aspergillus fumigatus* (ATCC-MYA-3626), *Rhizopus oryzae* (ATCC-9363), *Cunninghamella bertholletiae* (ATCC-42113), *Fusarium solani* (NBRC-5232), and a clinical isolate of *Cryptococcus neoformans*.

### 2.3 Fungal isolation, DNA extraction, and fungal identification

The fifteen fungal samples, collected from skin surfaces of the chest, abdomen, hands, and legs of cadavers, were initially cultured on Sabouraud Dextrose Agar (SDA) slants. These cultures were transported to Kumamoto University. Upon arrival, the fungi were subcultured onto SDA plates, and three representative colonies were selected for purification. After incubation at 30°C, total DNA was extracted from each of the resulting pure isolates.

DNA extraction protocol was previously described (Nakamura et al., 2024). In brief, fungal samples cultured on SDA were harvested, flash-frozen in liquid nitrogen, and then crushed using a BioMasher II (Nippi, Japan). Total DNA and RNA were subsequently extracted from the resulting material using TRIzol® reagent (Invitrogen, Thermo Fisher Scientific) according to the manufacturer’s protocol and stored at -80°C. For species identification, one of the following gene regions was amplified via PCR using 10–50 ng of template DNA: The targeted regions and their respective primer pairs were as follows: the Internal Transcribed Spacer (ITS) region using primers NS7F (5’-GAG GCA ATA ACA GGT CTG TGA TGC -3’) and ITS4R (5’-TCC TCC GCT TAT TGA TAT GC -3’); the β-tubulin gene using primers Bt2aF (5’-GGT AAC CAA ATC GGT GCT GCT TTC -3’) and Bt2bR (5’-ACC CTC AGT GTA GTG ACC CTT GGC -3’). All PCR amplifications were performed in a 40 µL reaction volume using KOD One® Polymerase (TOYOBO, Japan) in an Eppendorf® Mastercycler thermocycler.

### 2.4 Antifungal susceptibility testing

The minimum inhibitory concentrations (MICs) were determined by the broth microdilution method according to the Clinical and Laboratory Standards Institute (CLSI) document M38, 3rd edition (CLSI M38-Ed3, 2017), as previously described (Yoshinouchi et al., 2023). Serial twofold dilutions of each antifungal agent were prepared in MOPS-buffered RPMI 1640 medium. The fungal suspensions were then added to 96-well flat-bottom plates to achieve a final inoculum concentration of 4.0×10^3^ CFU/mL for filamentous fungi (*P. paneum, P. corylophilum, A. fumigatus, R. oryzae,* and *F. solani*), and 5.0×10^3^ CFU/mL for yeasts (*C. albicans* and *C. neoformans*). The plates were then incubated at 35°C for 48 to 72 hours. The MIC was defined as the lowest drug concentration (µg/L) that caused complete visual inhibition of fungal growth. The MIC_50_ and MIC_90_, the concentrations that inhibited 50% and 90% of fungal growth, respectively, were determined spectrophotometrically. This was achieved by measuring either the optical density at 530 nm (OD_530_) for fungal growth or the absorbance at 440 nm (OD_440_) following WST-1 cell viability staining. Data were taken using a FLUOstar Omega (BMG Labtech, Germany) and compared against a positive control (fungal growth without any drug) and a negative control (medium only).

### 2.5 Low-Temperature Growth Assay

To assess psychrotolerance, fungal suspensions of *P. paneum*, *P. corylophilum*, *A. fumigatus*, and *C. albicans* were adjusted to a final concentration of 5.0×10^3^ CFU/mL in 96-well flat-bottom plates and incubated at 4°C for 14 days. Fungal growth was monitored by measuring at OD_530_ on days 0, 1, 7, 10, and 14. On the final day of incubation, cell viability was also quantified using a WST-1 staining assay (OD_440_). All absorbance readings were performed with a FLUOstar Omega plate reader.

### 2.6 Fungal viability assay

Fungal viability was assessed using a WST-1 assay, as previously described (Nakamura et al., 2024). The WST-1 staining solution was prepared in 20 mM HEPES buffer (pH 7.2) and contained 5 mM WST-1 tetrazolium salt (Dojindo, Japan) and 0.2 mM 1-methoxy PMS (Dojindo, Japan) (Ishiyama et al., 1994). The solution was stored at -80°C until use. For the assay, 10 µL of the WST-1 solution was added to each well of the 96-well plate containing the fungal samples. The plate was then incubated at 35°C for 2–4 hours with shaking. Finally, the absorbance, corresponding to the amount of formazan produced by viable cells, was measured at 440 nm (OD_440_) using a FLUOstar Omega microplate reader.

### 2.7 Illustration

Illustrations accompanying the experimental procedure and explanations were created using BioRender (https://www.biorender.com/): Scientific Image and Illustration Software (Simplified Science Publishing, LLC [https://www.simplifiedsciencepublishing.com/]).

### 2.8 Drugs and disinfectants

Voriconazole (VRC), itraconazole (ITC), posaconazole (PSC), and terbinafine (TRB) were purchased from Tokyo Chemical Industry (Tokyo, Japan). Amphotericin B (AmB), 10% benzalkonium chloride, 20% chlorhexidine gluconate, and phenol were obtained from FUJIFILM Wako Pure Chemical Corporation (Osaka, Japan). Isavuconazole (ISC) was acquired from Selleck Chemicals (Houston, TX, USA), micafungin (MCF) from Cayman Chemical (Ann Arbor, MI, USA), and hypochlorous acid from Takasugi Seiyaku (Nara, Japan). Stock solutions of these compounds were prepared in dimethyl sulfoxide (DMSO) or other appropriate solvents at concentrations of 5–20 mM and stored at −80°C. For each assay, the stock solutions were thawed and diluted to the final working concentrations using MOPS-buffered RPMI medium.

### 2.9 DNA Sequences and GenBank Accession Numbers

The ITS sequences of *P. paneum* and *P. corylophilum* identified in this study were deposited in the DNA Data Bank of Japan (DDBJ) under accession number LC897326 and LC897327, respectively. The β-tubulin sequence of *P. corylophilum* were also deposited in the DDBJ, under accession number LC897497. DDBJ is linked to the International Nucleotide Sequence Database, INSD.

## 3. Results

### Isolation and molecular identification of fungi from cadaveric surfaces

Figure 1 illustrates the procedure for fungal identification from donated-cadaveric skin. Fungal growth was observed on the skin surface of cadavers at Institute of Science Tokyo. The cadavers were fixed by perfusion through the femoral artery using an 8% formalin solution. Before surgical dissection, they were preserved under refrigeration, wrapped in cloths saturated with 30% ethanol. To identify the fungi, samples were collected from skin areas showing fungal growth with varying colors and morphologies (Figure 1). The samples, initially cultured on Sabouraud Dextrose Agar (SDA) slants, were transported to Kumamoto University. There, they were subcultured onto SDA plates and incubated at 30°C to obtain pure isolates. Following the isolation and culture of the fungal colonies, total RNA and DNA were extracted. Species identification by sequencing of the Internal Transcribed Spacer (ITS) region and the β-tubulin gene revealed that 14 isolates were *Penicillium paneum* and one was *Penicillium corylophilum* (Sup. Table 1).

**Figure 1.**
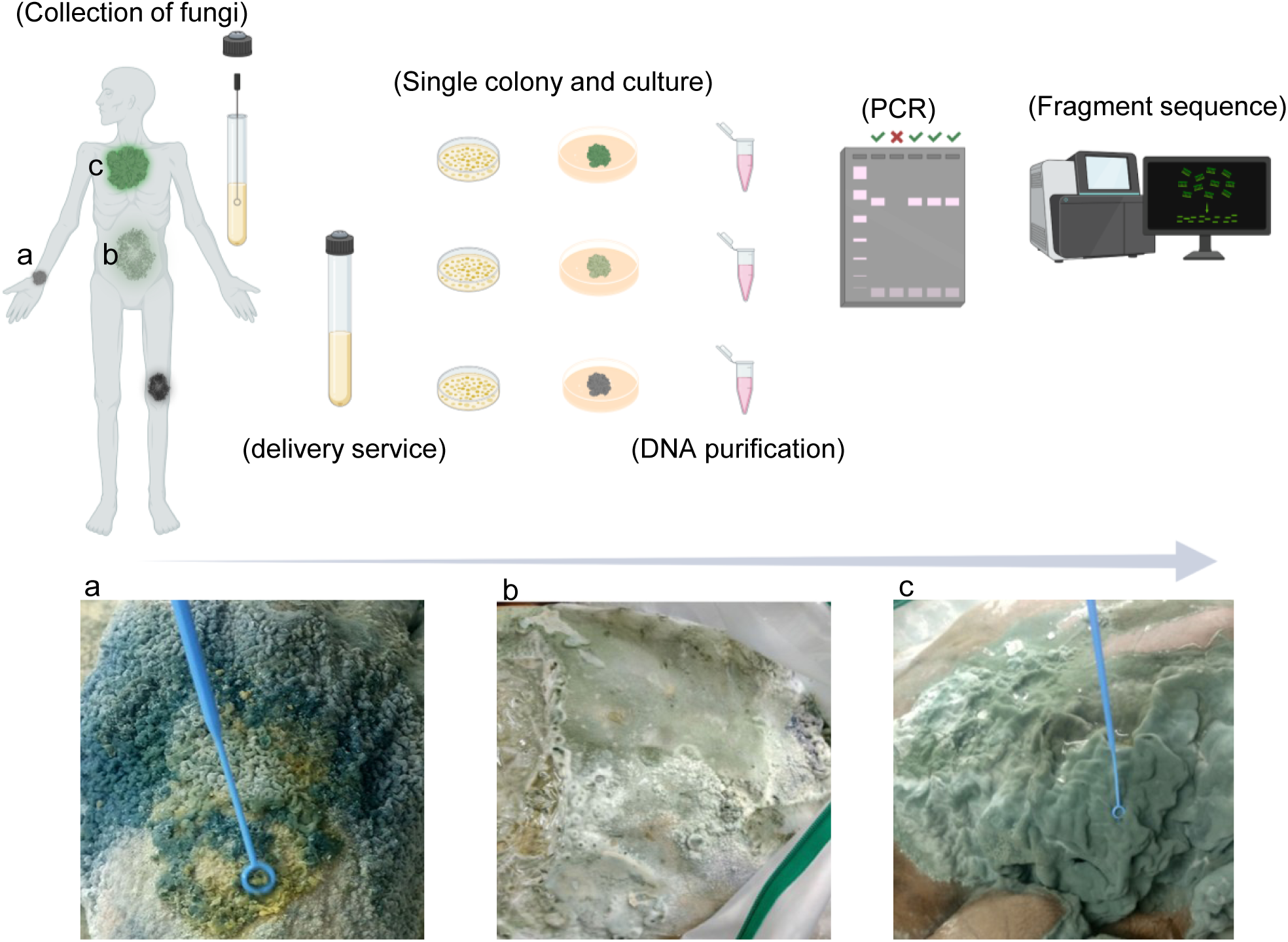
Workflow for the isolation and identification of fungi from cadaveric surfaces. Fungi exhibiting diverse morphologies and colors were observed on cutaneous surfaces of the chest, abdomen, hands, and legs of donated cadavers preserved under the institute of Science Tokyo method (a, b, c). Samples from these growths were collected, inoculated onto SDA slants, and transported to Kumamoto University. Pure isolates were obtained by subculturing the samples onto SDA plates and incubating them at 30°C. Total DNA and RNA was extracted from these isolates, and species were identified by Sanger sequencing of the ITS region and the β-tubulin gene.

### Morphological characterization of *P. paneum and P. corylophilum*

The macromorphological characteristics of both *P. paneum* and *P. corylophilum* in this study (Figure 2) were almost in agreement with the previous reports (Hocking et al., 1997; Visagie et al., 2014; O’brien et al., 2008). Colonies of *P. paneum* grown on SDA were circular and exhibited a velvety texture. They showed dense blue-green sporulation in the center, surrounded by a distinct white mycelial border. Light microscopy of specimens stained with Lactophenol-cotton-blue revealed complex, brush-like terverticillate conidiophore branching patterns characteristic of the species. SEM analysis further detailed these structures, showing phialides producing long chains of subglobose to round conidia. The overall structure appeared dense and compact (Figure 2A). On the other hand, *P. corylophilum* colony on SDA was white to pale grey, with a cotton-like texture. The colonies were raised and showed distinct radial furrows. Microscopic examination revealed biverticillate conidiophores with long, somewhat divergent metulae, resulting in a branching pattern less compact than that of *P. paneum*. SEM imaging confirmed this less compact hyphal arrangement and further showed that the conidia were orbicular, smooth-surfaced, and arranged in chains. (Figure 2B).

**Figure 2.**
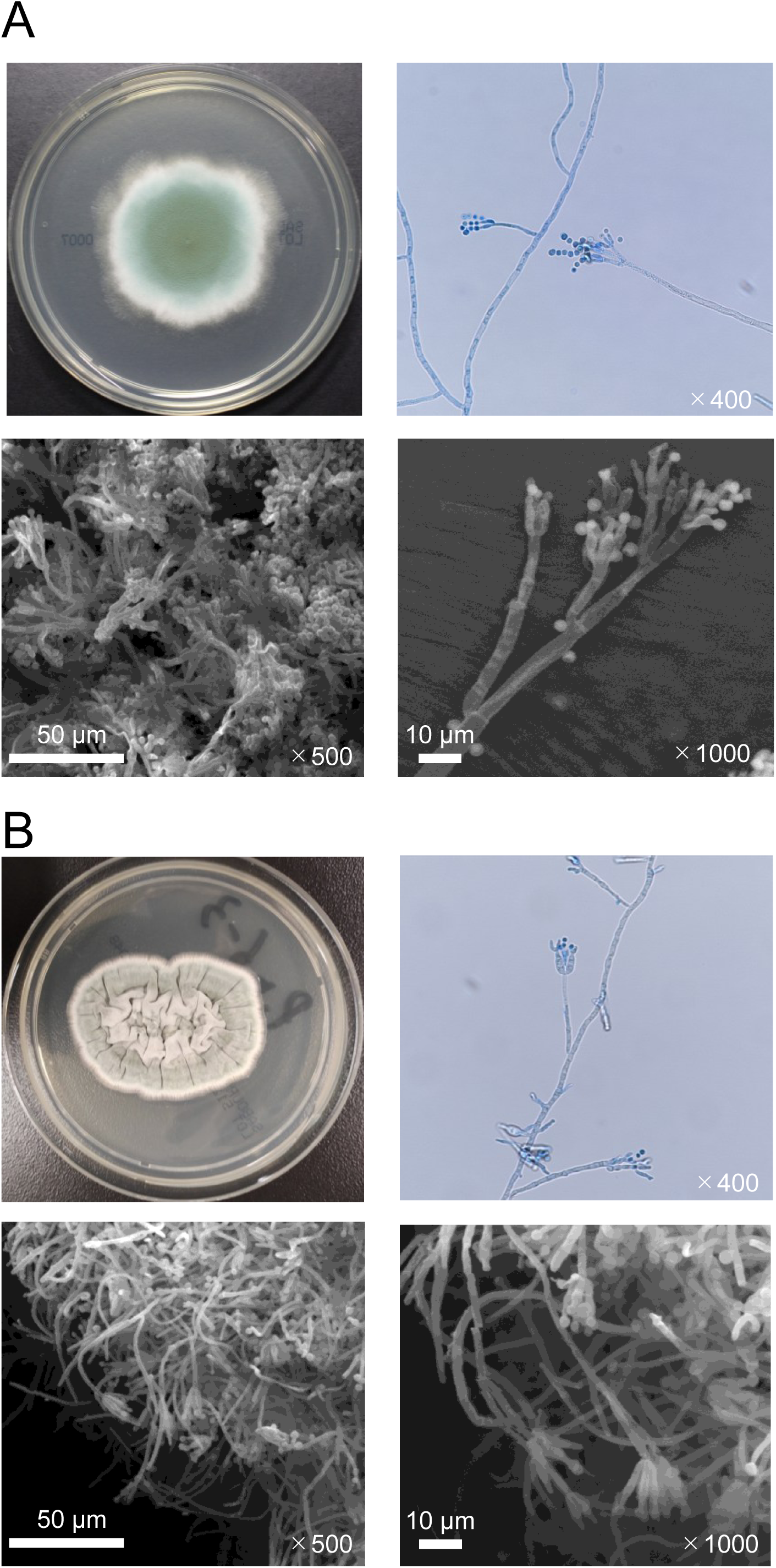
Fungal morphology of *P. paneum* and *P. corylophilum*. Morphology of (**A**) *P. paneum-1* and (**B**) *P. corylophilum-1*. Both isolates were cultured on SDA plates for 7 days at room temperature. Light micrographs show lactophenol cotton blue-stained specimens prepared by the slide culture method. Detailed structures were observed by scanning electron microscopy (SEM). Scale bars and magnifications are shown in each image.

### Psychrotolerance: growth capacity at 4°C

To investigate fungal growth of *P. paneum* and *P. corylophilum* at 4°C, two clones of *P. paneum-1* and *-2* and one clone of *P. corylophilum-1* were incubated in a 96-well plate for 14 days under refrigeration, along with *Aspergillus fumigatus* (Ben-Ami et al., 2010) and *Candida albicans* (Gow et al., 2017) as controls (Figure 3). Interestingly, *A. fumigatus* and *C. albicans* failed to grow, while a sharp increase in the proliferation of *P. paneum* and *P. corylophilum* was observed after 10 days (Figure 3A). Furthermore, *P. paneum*-*1*, *-2*, and *P. corylophilum-1* exhibited significant proliferation for 14 days at refrigeration temperatures using WST-1 assay staining for living cells compared with *A. fumigatus* and *C. albicans* (Figure 3B). These results suggest that both species are capable of growing at 4°C, which is consistent with the conditions for our cadaver preservation.

**Figure 3.**
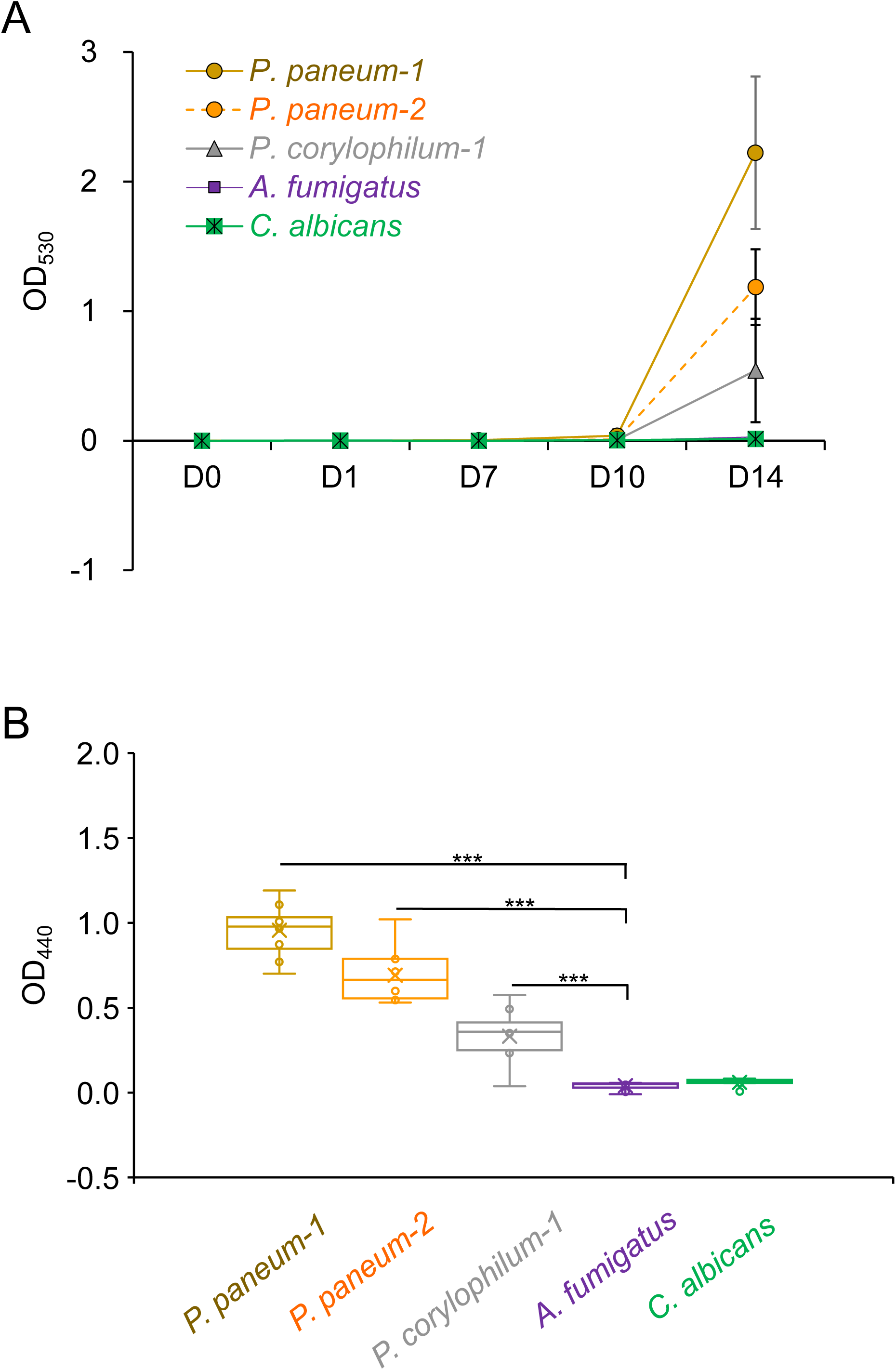
Psychrotolerance of *P. paneum* and *P. corylophilum* at 4°C. (**A**) Growth curves of *P. paneum-1*, *P. paneum-2*, *P. corylophilum-1*, *A. fumigatus*, and *C. albicans* incubated at 4°C were measured at 0, 1, 7, 10, and 14 days by OD_530_. (**B**) Cell viability of each strain after 14 days of incubation was determined by the WST-1 assay. Statistical comparisons between *Penicillium species* and control fungi groups were performed using Welch’s t-test (***, *P* < 0.0005).

### Ethanol tolerance and antifungal susceptibility profiles

Having established that these species can proliferate at refrigeration temperatures, we next investigated their tolerance to ethanol, another key component of our preservation environment. We compared the ethanol tolerance of *P. paneum* and *P. corylophilum* with that of several pathogenic fungi (*C. albicans*, *A. fumigatus*, *R. oryzae* (Gryganskyi et al., 2010), *C. bertholletiae* (Rickerts et al., 2000), *F. solani* (Zhang et al., 2006), and *C. neoformans* (Buchanan et al., 1998) stored in our laboratory. As shown in Table 1, *P. corylophilum* and *P. paneum* exhibited greater ethanol tolerance. This finding was supported by three methods: objective elevated MIC_50_ and MIC_90_ values were determined using optical density (OD_530_) and a WST-1 cell viability assay (OD_440_). These quantitative results were consistent with a subjective visual assessment of the MIC. The ethanol tolerance of the fungi was determined by their MIC values as follows: *P. paneum* (25%), *P. corylophilum* (25%), *A. fumigatus* (6.3%), *C. albicans* (3.1%), *R. oryzae* (1.6%), *C. bertholletiae* (1.6%), *F. solani* (0.8%), and *C. neoformans* (0.4%). In addition, the susceptibility of *P. paneum* and *P. corylophilum* to clinical antifungal drugs such as azoles (voriconazole, VRC; isavuconazole, ISC; itraconazole, ITC; posaconazole, PSC), terbinafine (TRB), amphotericin B (AmB), and micafungin (MCF) was determined, and the results are shown in Table 2 and Table 3. *P. paneum* was mildly more resistant to the tested antifungal drugs than *P. corylophilum*.

**Table 1.** Susceptibility of *P. paneum*, *P. corylophilum*, and control fungi to ethanol.

| fungi | OD<sub>530</sub><sup>b</sup> |  | OD<sub>440</sub> (WST-1)<sup>c</sup> |  | (%) |
| --- | --- | --- | --- | --- | --- |
|  | MIC<sub>50</sub> | MIC<sub>90</sub> | MIC<sub>50</sub> | MIC<sub>90</sub> | MIC<sup>a</sup> |
| *P. corylophilum* | 25.0 | 25.0 | 50.0 | 50.0 | 25.0 |
| *P. paneum* | 3.1 | 3.1 | 25.0 | 25.0 | 25.0 |
| *A. fumigatus* | 1.6 | 1.6 | 1.6 | 3.1 | 6.3 |
| *C. albicans* | 1.6 | 3.1 | 1.6 | 3.1 | 3.1 |
| *R. oryzae* | 0.8 | 3.1 | 3.1 | 3.1 | 1.6 |
| *C. bertholletiae* | 1.6 | 1.6 | 1.6 | 1.6 | 1.6 |
| *F. solani* | 0.8 | 0.8 | 0.8 | 1.6 | 0.8 |
| *C. neoformans* | 0.4 | 0.4 | 0.4 | 0.4 | 0.4 |

<sup>a</sup>The Minimum Inhibitory Concentration (MIC) was defined as the lowest ethanol concentration (%) that caused complete visual inhibition of fungal growth. <sup>b,c</sup>The MIC<sub>50</sub> and MIC<sub>90</sub> values represent the concentrations that inhibited 50% and 90% of fungal activity, respectively. Fungal growth was assessed by optical density at 530 nm (OD<sub>530</sub>), while cell viability was assessed by absorbance at 440 nm (OD<sub>440</sub>) after WST-1 staining. The data shown are representative of three independent experiments.

**Table 2.**
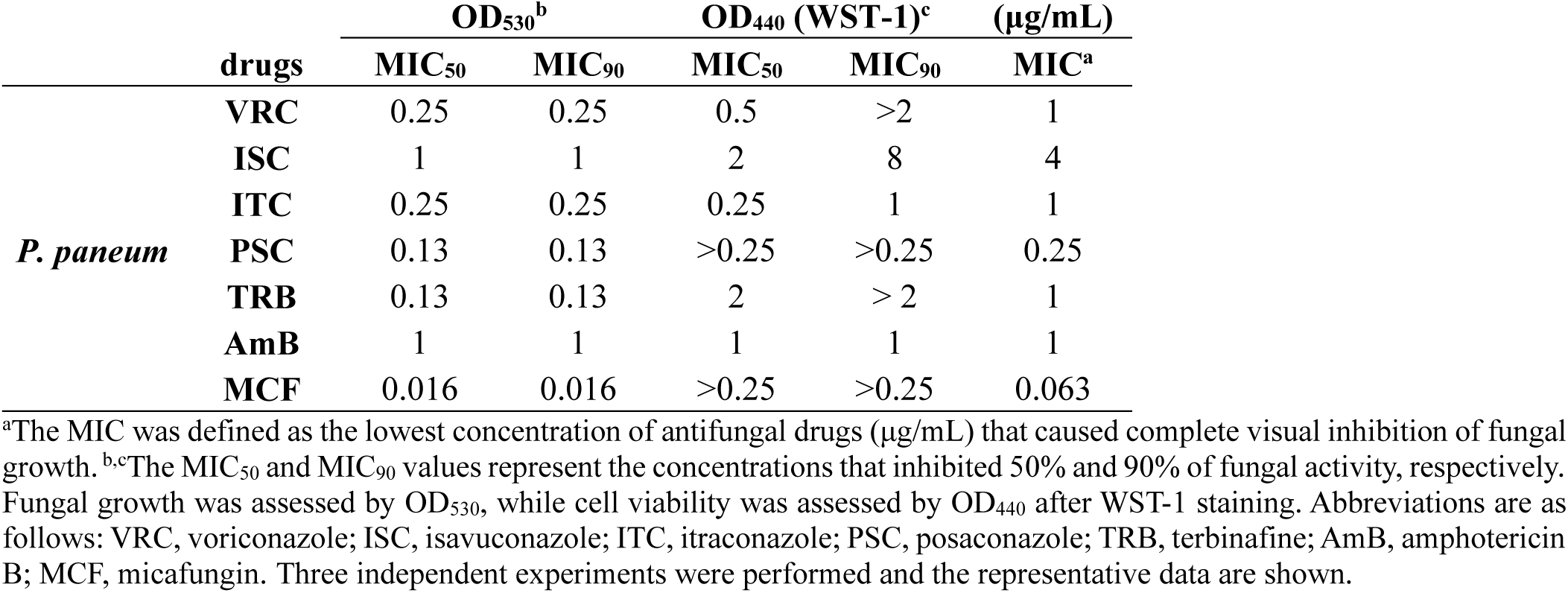
Susceptibility of *P. paneum* to clinical antifungal drugs.

**Table 3.**
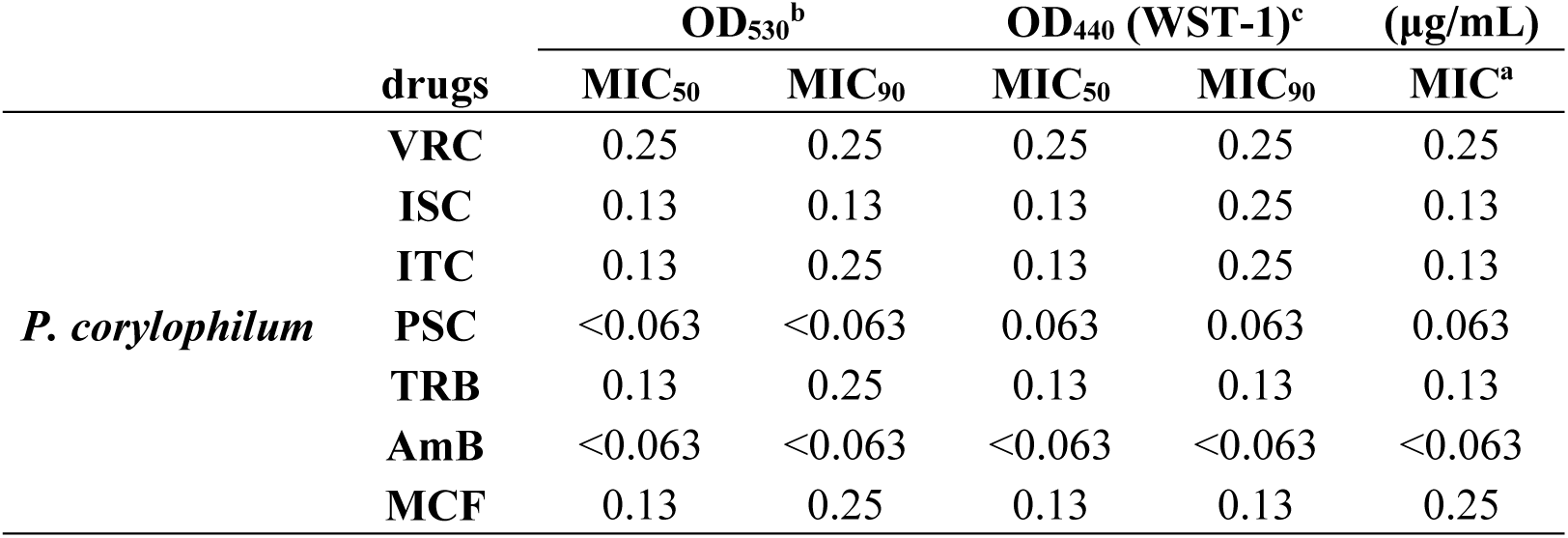
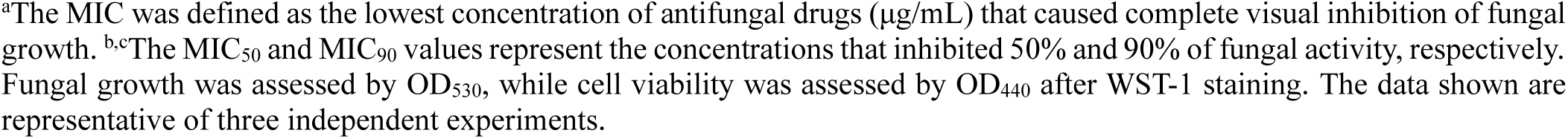
Susceptibility of *P. corylophilum* to clinical antifungal drugs.

### Efficacy of disinfectants against *Penicillium species*

To identify effective inhibitors of fungal growth on cadavers preserved under our conditions, we evaluated the efficacy of several disinfectants including benzalkonium chloride, chlorhexidine gluconate, phenol, and hypochlorous acid against *P. paneum* and *P. corylophilum* (Table 4). Benzalkonium chloride is used in disinfectant wipes and sprays for non-porous surfaces in hospitals, kitchens, and bathrooms (Merchel et al., 2019). Chlorhexidine gluconate is widely used by healthcare professionals as a hand scrub and for preparing a patient’s skin prior to an operation (DeRiso et al., 1996). Phenol is used as a surface disinfectant in laboratory settings, particularly for organic material contamination (DeBono et al., 1997). Hypochlorous acid is a high-level disinfectant used in hospitals and clinics to disinfect surfaces, medical equipment, and rooms (Block et al.,2020). Among the tested disinfectants, hypochlorous acid was the least effective, with high MICs against *P. paneum* and *P. corylophilum* in the range of 0.75–1.5% (Table 4). In contrast, the other agents were far more potent. Benzalkonium chloride was particularly effective, inhibiting the growth of both fungi at concentrations below 0.001%. Similarly, both chlorhexidine gluconate and phenol were highly effective, with MICs ranging from 0.001% to 0.004%. These results suggest that incorporating benzalkonium chloride, chlorhexidine gluconate, or phenol into cadaver preservation methods could prevent *P. paneum* and *P. corylophilum* from growing on the skin.

**Table 4.** Susceptibility of *P. paneum* and *P. corylophilum* to common disinfectants.

|  | *P. paneum* |  |  |  |  | *P. corylophilum* |  |  |  |  |
| --- | --- | --- | --- | --- | --- | --- | --- | --- | --- | --- |
|  | OD<sub>530</sub><sup>b</sup> |  | OD<sub>440</sub> (WST-1)<sup>c</sup> |  | (%) | OD<sub>530</sub><sup>b</sup> |  | OD<sub>440</sub> (WST-1)<sup>c</sup> |  | (%) |
| Disinfectants | MIC<sub>50</sub> | MIC<sub>90</sub> | MIC<sub>50</sub> | MIC<sub>90</sub> | MIC<sup>a</sup> | MIC<sub>50</sub> | MIC<sub>90</sub> | MIC<sub>50</sub> | MIC<sub>90</sub> | MIC<sup>a</sup> |
| **Benzalkonium chloride** | <0.001 | <0.001 | <0.001 | 0.002 | <0.001 | <0.001 | <0.001 | <0.001 | <0.001 | 0.002 |
| **Chlorhexidine gluconate** | 0.001 | 0.002 | 0.002 | 0.004 | 0.002 | 0.002 | 0.004 | 0.002 | 0.002 | 0.004 |
| **Phenol** | 0.008 | 0.016 | 0.016 | 0.016 | 0.016 | 0.004 | 0.004 | 0.002 | 0.004 | 0.004 |
| **Hypochlorous acid** | 1.5 | >1.5 | 0.75 | 1.5 | 1.5 | 0.75 | 1.5 | 0.75 | 1.5 | 1.5 |

<sup>a</sup>The MIC was defined as the lowest disinfectant concentration (%) that caused complete visual inhibition of fungal growth.
<sup>b,c</sup>The MIC<sub>50</sub> and MIC<sub>90</sub> values represent the concentrations that inhibited 50% and 90% of fungal activity, respectively.
Fungal growth was assessed by OD<sub>530</sub>, while cell viability was assessed by OD<sub>440</sub> after WST-1 staining. The data shown are representative of three independent experiments.

## 4. Discussion

This study identifies *P. paneum* and *P. corylophilum* as unique fungal contaminants capable of surviving under our cadaver preservation conditions: a low-formalin, ethanol-rich environment at 4°C. Their survival under these cadaveric conditions represents a novel finding, highlighting the ecological adaptability of these species and emphasizing the need for targeted antifungal countermeasures in anatomical preservation settings.

A key systematic review by Kwizera et al. highlights the issue of fungal contamination in cadavers (Kwizera et al., 2024). Their analysis of eight studies from Asia and the Caribbean found contamination on cadavers preserved in 5% to 14% formalin. The most prevalent contaminants identified across these studies were species of *Aspergillus*, *Penicillium*, and *Trichophyton*. Many microorganisms, including bacteria, yeast, and fungi, have enzymes that degrade formaldehyde into formate (Min et al., 2016). For example, formate oxidase has been purified from *Aspergillus nomius*, a formaldehyde-resistant soil fungus capable of completely consuming formaldehyde in media containing up to 0.45% (Kondo et al., 2002). It remains unclear whether the Penicillium species identified in our study possess similar enzymatic pathways. Furthermore, our preservation conditions differ significantly from other standard techniques, such as the Thiel method, which utilizes a different chemical composition.

On the other hand, the findings of this study regarding remarkable tolerance of *P. paneum* and *P. corylophilum* to both chemical (ethanol) and physical (4°C) stressors may have broader implications. For example, this tolerance is particularly relevant to the related field of Forensic Mycology, which has traditionally focused on fungi found on cadavers as important evidence (Martínez-Ramírez et al., 2013). Forensic Mycology helps estimate the Post-mortem Interval (PMI), detect the translocation of remains, and identify the original crime scene (Spychała et al., 2024). Understanding which fungal species can survive extreme or unusual conditions, as demonstrated here, could potentially refine these forensic analyses in atypical scenarios.

Regarding the established characteristics of these species, *P. paneum* is a filamentous fungus known as a significant spoilage organism in food and animal feed, particularly silage. The primary health concern is not direct infection, but the threat to livestock from its potent mycotoxins, including patulin and roquefortine C. The fungus is reported to be highly adapted to these environments, exhibiting tolerance to low pH, low temperatures, and high-CO₂/low-O₂ atmospheres. Taxonomically related to *P. roqueforti*, this species is also recognized as a plant pathogen and an agent of biodeterioration (Visagie et al., 2014, O’brien et al., 2008). *P. corylophilum* is a ubiquitous environmental fungus, commonly isolated from soil and damp indoor environments. Clinical significance is chiefly derived from its role as a powerful aeroallergen and a producer of mycotoxins, including the nephrotoxic agent citrinin. *P. corylophilum*’s inability to grow effectively at 37°C is believed to account for the rarity of direct infections. A key physiological characteristic of this species is its xerophilic nature, enabling it to colonize substrates with low water activity, making it a frequent indoor contaminant (Hocking et al., 1997).

Antifungal susceptibility testing revealed high sensitivity to benzalkonium chloride, chlorhexidine gluconate, and phenol, suggesting that incorporation of these agents into ethanol-based preservation fluids may effectively suppress fungal colonization. This practical strategy aligns microbiological understanding with occupational safety in anatomy education facilities.

This study has several limitations. First, we examined a limited number of cadavers from a single institution. Second, we did not investigate the enzymatic mechanisms underlying formaldehyde degradation in these species. Third, the long-term efficacy of the proposed disinfectants in actual preservation conditions requires validation through longitudinal studies

In conclusion, this study demonstrates that a thorough understanding of a fungal ecology is fundamental to effectively managing and resolving any contamination.

## CRediT authorship contribution statement

**Tatsuya Yoshinouchi:** Investigation, Formal analysis, Validation. **Tomofumi Nakamura:** Conceptualization, Writing - original draft, Visualization, Investigation, Formal analysis, Visualization, Project administration. **Masahiro Manabe:** Investigation. **Daisuke Mori**: Writing - Review & Editing, Supervision. **Jun-ichirou Yasunaga:** Writing - Review & Editing, Supervision. **Yasuhito Tanaka:** Writing - Review & Editing, Supervision. **Joe Iwanaga:** Writing - review & editing, Supervision. **Norio Kitagawa:** Conceptualization, Writing - review & editing, Resources, Project administration.

## Acknowledgments

We also thank Mayu Okumura, Hirotomo Nakata, and Mitsuhiro Uchiba for discussion of the data. The authors sincerely thank those who donated their bodies to science so that anatomical research could be performed. Results from such research can potentially increase mankind’s overall knowledge that can then improve patient care. Therefore, these donors and their families deserve our highest gratitude. The authors hereby confirm that every effort was made to comply with all local and international ethical guidelines and laws concerning the use of human cadaveric donors in anatomical research (Iwanaga et al., 2021; Iwanaga et al., 2022; Iwanaga et al., 2025).

## Appendix. Supplementary data

**Sup. Table.1** The sequences ITS region and β-tubulin gene of collected fungi

## Declaration of AI and AI-assistant technology in the writing process

During the preparation of this work, the authors used Gemini Pro to correct spelling and grammatical errors and to improve English readability. After using this tool/service, the authors reviewed and edited the content as needed and take full responsibility for the content of the published article.

